# A temporally resolved transcriptome for developing “Keller” explants of the *Xenopus laevis* dorsal marginal zone

**DOI:** 10.1101/2020.09.22.308312

**Authors:** Anneke D. Kakebeen, Robert Huebner, Asako Shindo, Kujin Kwon, Taejoon Kwon, Andrea E. Wills, John B. Wallingford

## Abstract

Explanted tissues from vertebrate embryos reliably develop in culture and have provided essential paradigms for understanding embryogenesis, from early embryological investigations of induction, to the extensive study of *Xenopus* animal caps, to the current studies of mammalian gastruloids. Cultured explants of the *Xenopus* dorsal marginal zone (“Keller” explants) serve as a central paradigm for studies of convergent extension cell movements, yet we know little about the global patterns of gene expression in these explants. In an effort to more thoroughly develop this important model system, we provide here a time-resolved bulk transcriptome for developing Keller explants.

## Introduction

The culture of explanted tissues of early embryos has been a key tool for understanding vertebrate development for nearly a century, since Johannes Holtfreter first defined a medium capable of supporting the development of explanted vertebrate tissues. His landmark studies described the developmental potential of nearly every region of early amphibian embryos (translated in: (Hamburger, 1996a; Hamburger, 1996b)). Though amphibia are especially amenable to such explantation, similar early investigations were also made in the fish (Oppenheimer, 1936). Later in the 20th century, explanted amphibian tissues from *Xenopus* played a key role in early studies of the molecular basis of vertebrate development (Fraser and Harland, 2000; Heasman, 2006).

More recently, there has been renewed interest in the study of early embryo explants from zebrafish (Fulton et al., 2020; Schauer et al., 2020; Williams and Solnica-Krezel, 2020; Xu et al., 2014), which has prompted comparisons to mammalian gastruloids, in which pluripotent stem cells are directed toward early embryonic fates (Simunovic and Brivanlou, 2017). Indeed, explants of the fish blastoderm have even been referred to as “pescoids” and the hugely tractable explants of pluripotent blastula ectoderm from *Xenopus* (so-called “animal caps”) as “Xenoids” (Moris et al., 2020). Whether these new entries into the developmental biology lexicon will be widely adopted remains to be seen. Regardless, the recent interest in “oids” reminds us that among the more deeply studied tissue explant systems is the dorsal marginal zone (DMZ) of the amphibian gastrula.

Taking its name from its origin at the margin of the animal and vegetal halves on the future dorsal side of the embryo, the DMZ tissue is fated to become notochord, somite, and central nervous system (Dale and Slack, 1987; Keller, 1975; Keller, 1976; Moody, 1987). However, the tissue is more famously known as the “Spemann-Mangold Organizer,” and if the DMZ of the early gastrula is transplanted to the ventral marginal zone, it will re-pattern the surrounding host tissue, causing it to take on dorsal cell fates (Martinez Arias and Steventon, 2018; Sander and Faessler, 2001).

Interestingly, in his earliest studies of explants, Holtfreter noted that the DMZ will elongate (Hamburger, 1996a), and both he and Schechtman studied this explanted tissue extensively to show that such elongation was an autonomous property (Holtfreter, 1943; Holtfreter, 1944; Schectman, 1942). Decades later, Ray Keller and his lab’s pioneering work on convergent extension movements established *Xenopus* DMZ explants (aka “Keller” explants) as a key paradigm for studies of vertebrate morphogenesis (Keller and Danilchik, 1988; Keller and

Hardin, 1987; Shih and Keller, 1992a; Shih and Keller, 1992b; Wilson and Keller, 1991). Since then, the DMZ explant has been extensively interrogated as a model for diverse aspects of convergent extension, ranging from the molecular to the mechanical (e.g. (Habas et al., 2003; Kim and Davidson, 2011; Moore et al., 1995; Shindo and Wallingford, 2014; Shook et al., 2018; Skoglund et al., 2008; Tahinci and Symes, 2003; Wallingford et al., 2000)).

The DMZ is thus an especially interesting explant: It is a key source of inductive signals, it undergoes differentiation into several distinct tissues, and it undergoes a robust, autonomous process of morphogenesis. Despite the important role these explants continue to play in developmental biology, we still know relatively little about their transcriptional dynamics. For example, only a small number of genes have been examined by *in situ* hybridization in DMZ explants, (e.g. (Davidson et al., 2004; Doniach et al., 1992; Poznanski and Keller, 1997; Ruiz i Altaba, 1992; Saint-Jeannet et al., 1994; Sater et al., 1993)). Moreover, though we and others have reported the large-scale transcriptome for the early gastrula DMZ of *Xenopus* (Ding et al., 2017; Popov et al., 2017), these datasets are temporally limited.

In an effort to improve the utility of this explant system, we extend previous work here by providing a time-resolved bulk transcriptome for *Xenopus* DMZ “Keller” explants from early gastrulation through late neurulation. We find that these explants develop the full extent of A/P and D/V pattern, though certain elements of this pattern appear not to be maintained at later stages. We further find that, consistent with previous reports, close attention should be paid to the L and S homeoalleles when selecting molecular markers. Finally, we find a surprising lack of syn-expression of genes implicated in the control of convergent extension.

## Results and Discussion

To aid in understanding how cells in the DMZ acquire distinct fates and initiate specific behaviors over developmental time, we sought to define the dynamic transcriptome of this explant. We manually dissected DMZ explants at stage 10.5 and cultured them in 1x Steinberg’s solution (Figure 1A, B). We then collected 5 DMZs at stages from 11-18 (Nieuwkoop and Faber, 1967), and prepared RNA-Seq libraries in duplicate from the explants at each stage (Figure 1C). To provide outgroups, we also prepared RNA-Seq libraries in duplicate from five ventral marginal zone (VMZ) explants isolated at stage 10.5 and cultured to stage 11, as well as 20 animal cap explants isolated at stage 8 and cultured to stage 11 (Figure 1A).

**Figure 1.**
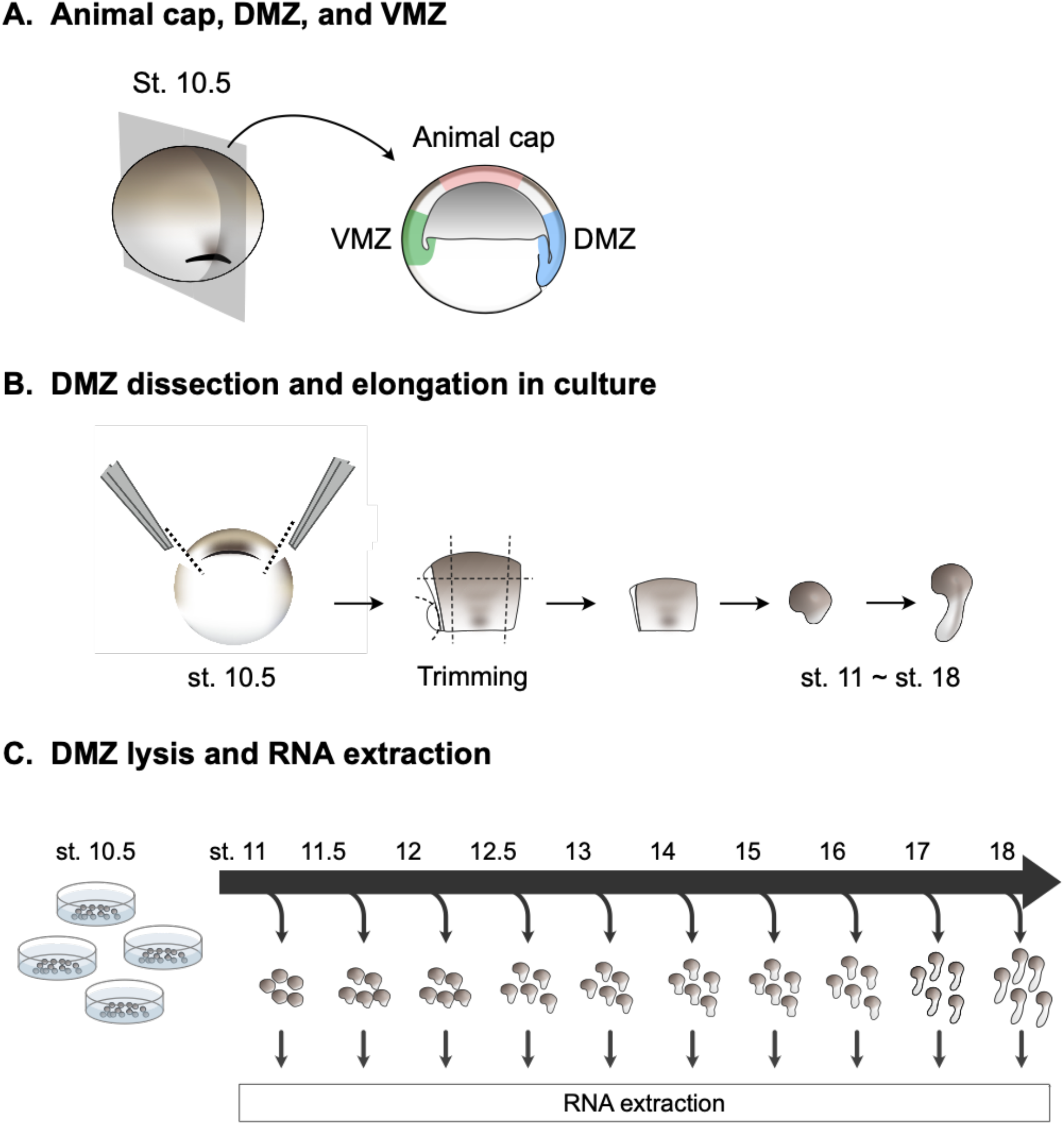
Dissections, culturing, and preparation of explant RNA-Seq libraries. A) Diagram identifying where cuts were made for dorsal marginal zones (DMZ), ventral marginal zones (VMZ), and animal caps. B) Diagram showing flow of dissection and culturing until RNA extraction. C) Diagram showing DMZ explant culture and RNA collection timeline.

To assess the quality of our dissections, we first compared the set of genes enriched in the DMZ over the VMZ at stage 11 in our dataset to our previous similar dataset, as well as to another independently generated dataset (Ding et al., 2017; Popov et al., 2017). We were surprised to observe relatively modest overlap, but this was the consistent trend for all three studies (Figure 2A). Critically, however, despite the overall modest overlap, the set of genes enriched across all three studies included many of the most well-known dorsally-restricted transcripts, including *noggin, chordin, follistatin, goosecoid*, and *nodal*, all of which play key roles in Spemann-Mangold Organizer specification or function (Figure 2B). Others, such as *tmem150b, lrig3* and *sds* are known to be expressed dorsally in the gastrula, though their function in this context is unclear (Bourdelas et al., 2004; Sinner et al., 2006; Zhao et al., 2008).

**Figure 2.**
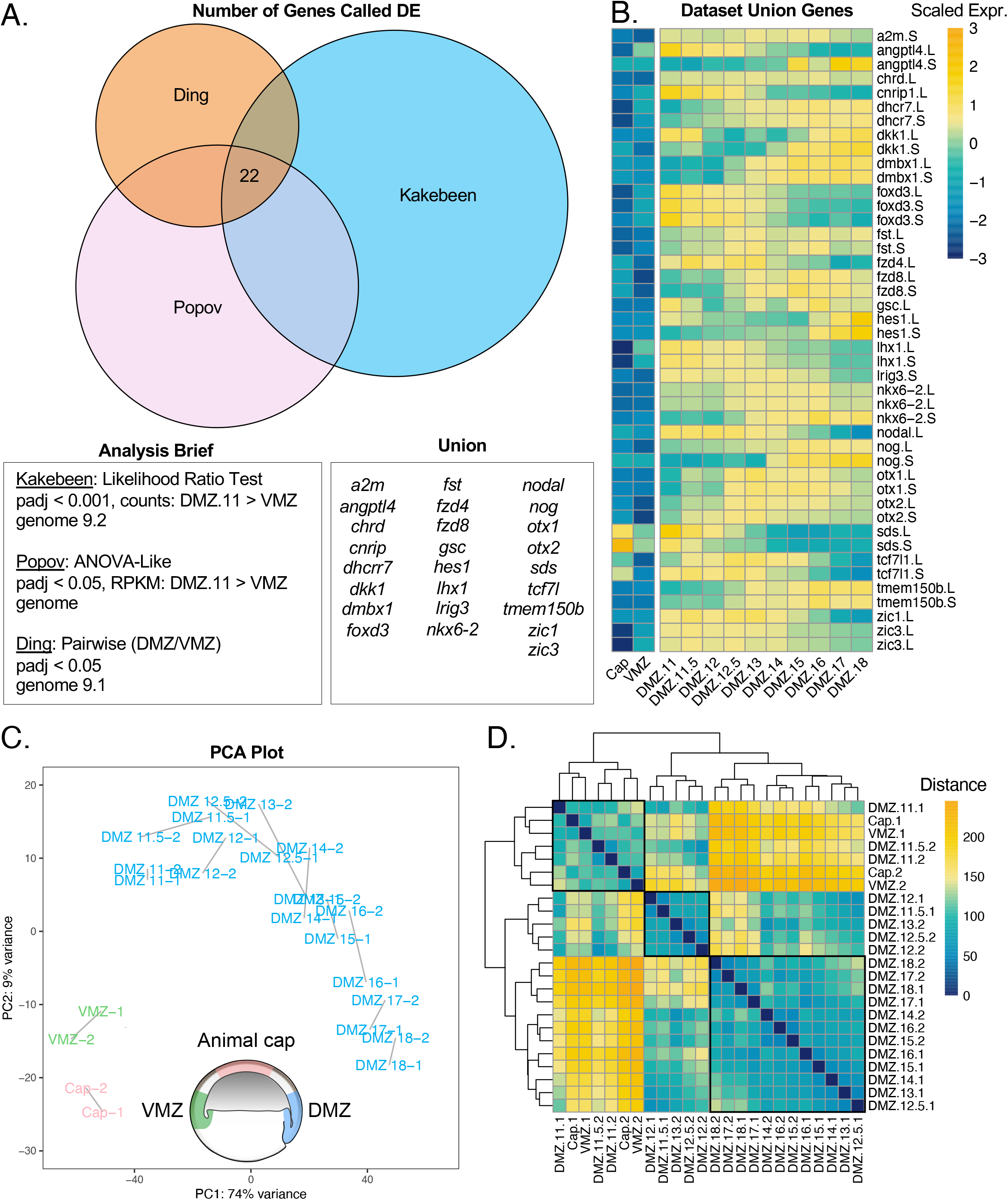
Cultured DMZs can be temporally differentiated by their global transcriptomes. A) Venn diagram comparing genes enriched in the DMZ with respect to the VMZ in three independent studies. Differential expression analysis for each data set are reported followed by the genes representing the union of all three data sets. B) Heatmap showing gene expression of genes captured by the union of all three data sets. Color represents expression of gene scaled by row. C) PCA plot clustering all samples. Blue samples are cultured DMZ, green samples are cultured VMZ, and pink samples are cultured animal caps. D) Correlation heatmap demonstrating the calculated distance between samples. Hierarchical clustering was performed by Pheatmap.

We reasoned that the modest overlap between the three studies relates predominantly to technical variation, not biological. Therefore, to test the variation in technical approaches, we compared genes called as enriched in our own datasets for DMZ stage 11 over VMZ stage 11 using three different analysis methods. Two of the analyses were performed with DESeq2 downstream of alignment and quantification with kallisto (Bray et al., 2016) Supp. Figure 1A-LRT/Wald), and one analysis was performed with limma (Ritchie et al., 2015) downstream of alignment with STAR (Dobin et al., 2013)(Supp Fig. 1A-Limma). We found that the genes called by the likelihood ratio test (LRT) and Wald test largely overlapped, whereas the overlaps between the genes called by limma and LRT or Wald overlapped less (Supp. Figure. 1A). The union of all three comparisons returned 100 genes which made up 14% of the LRT genes, 30% of the Wald genes, and 24% of the limma genes. The genes called in the union were all more enriched across all DMZ samples than the VMZ or animal caps (Supp. Figure 1B). Although we returned DMZ-enriched genes with high confidence, it is clear that the analysis pipeline used can influence the transcripts resolved. Despite the overall variability between the datasets, the strong enrichment of core marker genes made us confident that our dissections were effective.

We next used principal component analysis (PCA) to examine the quality of our samples and to identify characteristics that differentiate the samples (Figure 2C). This analysis demonstrated the strong reproducibility of our biological duplicates, as expected, and also identified the animal cap and the VMZ as outgroups, again as expected (Figure 2C). We also found that PC1, which accounts for 74% of the variance between samples, corresponds to the temporal ordering of samples (Figure 2C). Visualizing a distance matrix between the samples also further showed that samples are grouped temporally into three main clusters (Figure 2D). The first cluster groups animal cap, VMZ, and DMZ stage 11-11.5; the second groups DMZ stage 11.5-13; and the third groups DMZ stage 13-18. Although these are the largest clusters, there are small subclusters that group with neighboring timepoints. Together, these data suggest aged DMZ samples can be temporally resolved by their global gene expression.

Our goal was to identify dynamic patterns of gene expression over developmental time in Keller explants, so we next clustered differentially expressed genes across all DMZ samples. We identified genes that were differentially expressed between samples in our dataset using DESeq2’s likelihood ratio test (LRT) to statistically identify genes that were differentially expressed between any two samples (Love et al., 2014). We further restricted the list of differentially expressed genes by filtering our results for an FDR value of less than 0.001 and recovered 13,913 genes. To identify temporal patterns of gene expression, we applied k-means clustering to the set of all differentially expressed genes in DMZ samples. All transcripts recovered, differentially expressed transcripts, and cluster designations can be found in Supplemental Table 1.

Our k-means clustering analysis resulted in five clusters of syn-expression which described three general trends in the data (Figure 3A): First, slightly less than half of genes displayed a fairly consistent reduction in expression over time, with the onset of reduced expression lagging somewhat in cluster 2 compared to cluster 1. Second, slightly less than half of genes displayed roughly the converse expression pattern, showing a smooth, consistent increase in expression over time, with cluster 4 reaching a peak somewhat earlier than Cluster 5. Finally, a comparatively small set of genes displayed a peak of expression at mid-neurulation, with lower expression at early and late stages; all of these genes clustered together in Cluster 3.

**Figure 3.**
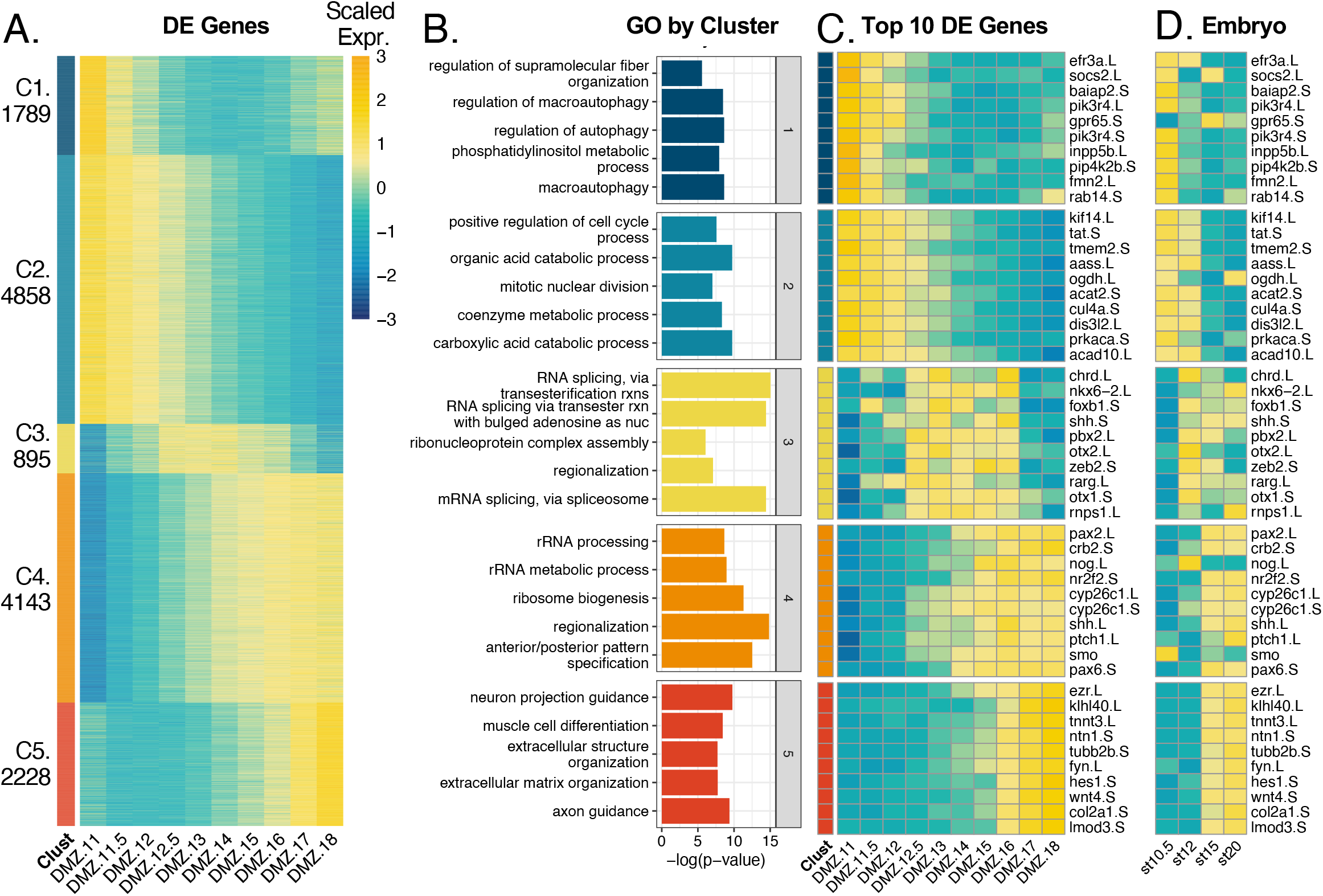
Cultured DMZs cluster to 5 k-means clusters. A) Heatmap of all differentially expressed genes clustered to 5 k-means clusters. B) Gene ontology analysis results from GO analysis on all genes per cluster. C) Heatmap of top 10 genes by FDR from each cluster. Color in heatmaps represent expression of gene scaled by row D) Heatmap of the same genes in C, plotted from published data in the whole embryo at complimentary developmental timepoints.

We used gProfiler2 (Raudvere et al., 2019) to define a ranked list of genes that represent each cluster by gene ontology analysis (Figure 3B). Curiously, we found that cluster 1 genes were most enriched for terms relating to autophagy and lipid metabolism. The autophagy terms included several highly expressed genes in the cluster such as *efr3a, socs2, baiap2, pik3r4, gpr65, inpp5b, pip4k2b, fmn2, and rab14* (Figure 3C). Interestingly, this cluster also includes the ubiquitin ligase *nedd4L*, which is required not only for autophagy, but also for specification and morphogenesis of the DMZ (Nazio et al., 2016; Zhang et al., 2014).

Cluster 2 genes were enriched for metabolic processes and regulation of cell cycle processes. The most significant, differentially expressed genes from these terms were *kif14, tat, tmem2, aass, ogdh, acat2, cul4a, dis3l2, prkaca, and acad10* (Figure 3C). Cluster 3 genes were enriched for processes related to mRNA splicing and regionalization. The most significant differentially expressed genes in this cluster (*chrd, nkx6-2, foxb1, shh, pbx2, otx2, zeb, rarg, otx1, and rnps1*) were responsible for driving the term “regionalization” (Figure 3C). The timing of high expression in this cluster coincides well with when neural tube regionalization typically occurs in the embryo. Clusters 3 and 4 have an overlap in stages with high gene expression, thus they call similar types of GO terms. Genes from cluster 4 are enriched for rRNA terms, regionalization, and anterior/posterior specification. Genes such as *pax2, crb2, nog, nr2f2, cyp26c, shh, ptch, smo, and pax6* were differentially expressed and enriched for the regionalization and specification terms (Figure 3C).

Although clusters 4 and 5 have similar expression patterns, cluster 4 genes turn on expression around stage 13, thus coinciding with stages still tasked with regionalization and specification. Cluster 5 has a tighter domain of expression from stage 16-18, resulting in a more divergent set of GO terms relating to cell differentiation specifically for neurons and muscle cells. Differentially expressed genes such as *ezr, ntn1, tubb2b*, and *fyn* contribute to neural terms whereas *klhl40, tnnt3, hes1*, and *wnt4* contribute to muscle differentiation (Figure 3C). Perhaps most interestingly, the genes most well-known for controlling key developmental events in *Xenopus (chordin, noggin, shh, otx2, pax2*) were found in clusters 3 and 4, showing peak expression at later gastrula stages. Conversely, the functions of genes in clusters 1 and 2, with peak expression early in gastrulation, are far less studied in the context of *Xenopus* development.

To compare the development of isolated DMZs to their normal development *in vivo*, we compared the temporal patterns of expression for the top ten genes in each of the five clusters with temporal patterns of expression from published RNA-Seq for these genes in the whole embryo (Session et al., 2016). We found remarkable concordance of temporal trends in gene expression, though with some notable exceptions (Figure 3D). For example, while the expression of Shh, Nkx6.2 and Foxb1 remains robust until well after stage 18 in whole embryos, all three genes displayed strongly reduced expression at late neurula stages in DMZ explants (Fig. 3C, D). Because all three genes are implicated in D/V patterning of the neural tube (Dichmann and Harland, 2011; Peyrot et al., 2011; Takebayashi-Suzuki et al., 2011; Zhao et al., 2007), this result may suggest that establishment of this D/V pattern can proceed in isolated DMZs but maintenance of the pattern requires an external signal from an embryonic tissue not present in the explant (the endoderm normally underlying the midline, for example).

We next considered that decades of studies of early axial patterning in *Xenopus* have relied on the use of a common, relatively small set of molecular markers for dorsoventral patterning of the gastrula, neuroectodermal patterning, and dorsoventral patterning of trunk mesoderm (e.g. (Elkouby and Frank, 2010; Essex et al., 1993; Heasman, 2006; Knecht and Harland, 1997; Sasai and De Robertis, 1997)). We therefore examined the expression of these “canonical” molecular markers in our dataset (Figure 4). All of the commonly used markers of early dorsoventral patterning in the gastrula were expressed as expected: BMP antagonists *noggin* and *chordin* in the DMZ, *bambi, vent2*, and *szl* in the VMZ. Interestingly, however, all of the ventral markers tested were also strongly expressed in our animal cap samples (Figure 4). This result does not seem to be an artifact of the animal cap dissection, as the canonical pan-mesodermal marker *xbra/t* was robustly present in both the DMZ and VMZ samples, but not in caps. Conversely, the animal cap samples were enriched in epidermal keratins compared to VMZs, as expected.

**Figure 4.**
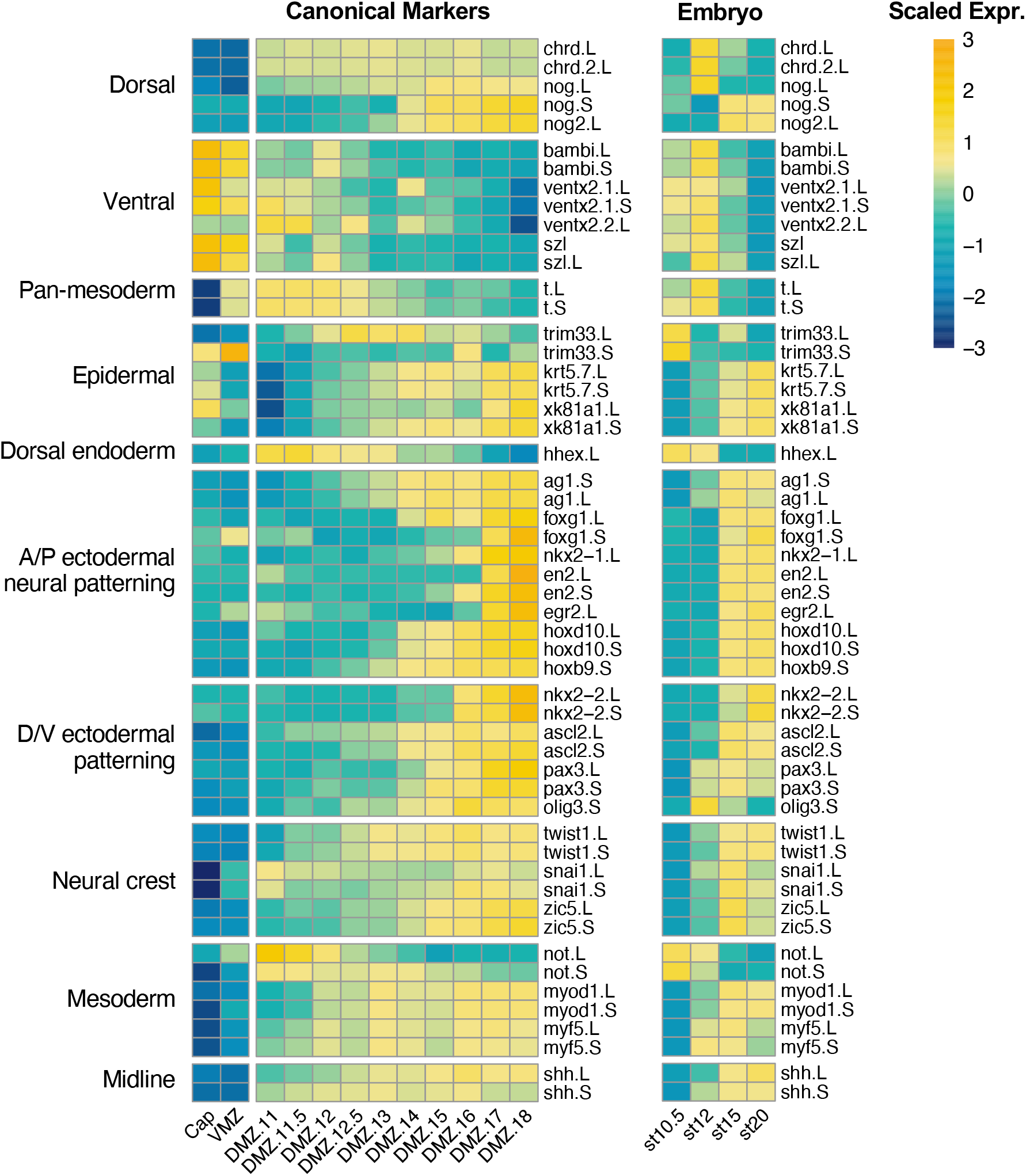
Canonical markers used to study development of the Xenopus embryo. Heatmaps showing gene expression of canonical markers of development in (left) explants and (right) whole embryos (Session 2016). Color represents expression of gene scaled by row.

Also of interest, our data show that Keller explants recapitulate the full extent of anteroposterior pattern of the ectoderm. They developed robust expression of anterior ectodermal markers such as the cement gland marker *xag1 (ag1*), the forebrain markers *foxg1/xbf1* and *nkx2.1;* the mid- and hindbrain markers *en2* and *egr2/krox20*, and the hindbrain/spinal cord markers *hoxd10* and *hoxb9* (Figure 4). Perhaps more surprisingly, these explants also display robust expression of neural markers along the full length of the dorsoventral axis, from *nkx2.2* in the floorplate, to the interneuron marker *ascl2/xash3*, the dorsal neural markers *pax3* and *olig3*, and several neural crest markers (e.g. *twist, sox10*). Thus, the full extent of AP and DV pattern is acquired in isolated Keller explants, providing a fuller picture of the efficacy of planar neural induction in *Xenopus* (e.g. (Doniach et al., 1992; Poznanski and Keller, 1997; Ruiz i Altaba, 1992; Sater et al., 1993)).

From a practical standpoint, this dataset will be useful for the design of future studies. First, the data provide an improved view of the temporal pattern of expression for these markers (e.g. *xnot* provides good marking of the notochord early, but *noggin* will be better at later stages). Moreover, *Xenopus laevis* has an allotetraploid genome (Session et al., 2016) yet the L and S form of each gene were until recently rarely considered in the selection of markers. Our data makes clear that this significant factor must not be overlooked: For example, both the L and S allele of the mesodermal marker *xbra* (*t*) are clearly absent from animal caps but are strongly expressed in both VMZ and DMZ and have similar temporal expression patterns (Figure 4). By contrast the S allele of the commonly used organizer marker *noggin* is only very weakly expressed and starts to be more strongly expressed in later stages. Differences in temporal gene expression also are informative for choosing between alleles of the key ectodermal regulator *ectodermin/trim33*. While both genes are expressed in animal caps with similar expression levels, gene expression increases in the DMZ samples for L and decreases in the DMZ samples, suggesting *trim33.S* may be the most appropriate marker to use to discriminate between epidermis and mesoderm.

The divergence in expression levels and temporal expression patterns between L and S homoeologues of some markers prompted us to ask how well L and S gene expression and temporal patterns of expression are correlated genome-wide. To this end we first asked whether both homeoalleles are recovered at similar rates throughout our data. Overall, we found two-thirds of the transcripts recovered could be confidently assigned to the L or S genome. Genes with both L and S transcripts were only recovered for 40% of the genes in our dataset (Figure 5A). We only recovered L transcripts for a comparable proportion of genes, and only S transcripts from about 19% of genes. If we consider only differentially expressed genes, the proportion of genes with both differentially expressed L and S homoeologues increased to 51%, whereas the percent of L or S only genes decreased (Figure 5B). For genes where both L and S transcripts were expressed in our data set, we next asked if the expression dynamics of L and S homoeologues were generally similar over time. We therefore calculated a Pearson correlation coefficient for the expression of all L vs S pairs across all tissue types and stages. L and S homoeologous pairs for the same gene are significantly better correlated than randomly paired L and S of different genes (Figure 5, C versus D). We then assigned correlation coefficients for homoeologues of each gene over the DMZ timecourse and visualized these by a histogram (Figure 5E). We found that correlation coefficients fell between 0.75 and 1 for most genes, but there are many with coefficients less than 0.75, or even less than 0.5 (Figure 5E). Thus, high (or low) expression of the L homoeologue is only somewhat predictive of similar expression for the S homoeologue.

**Figure 5.**
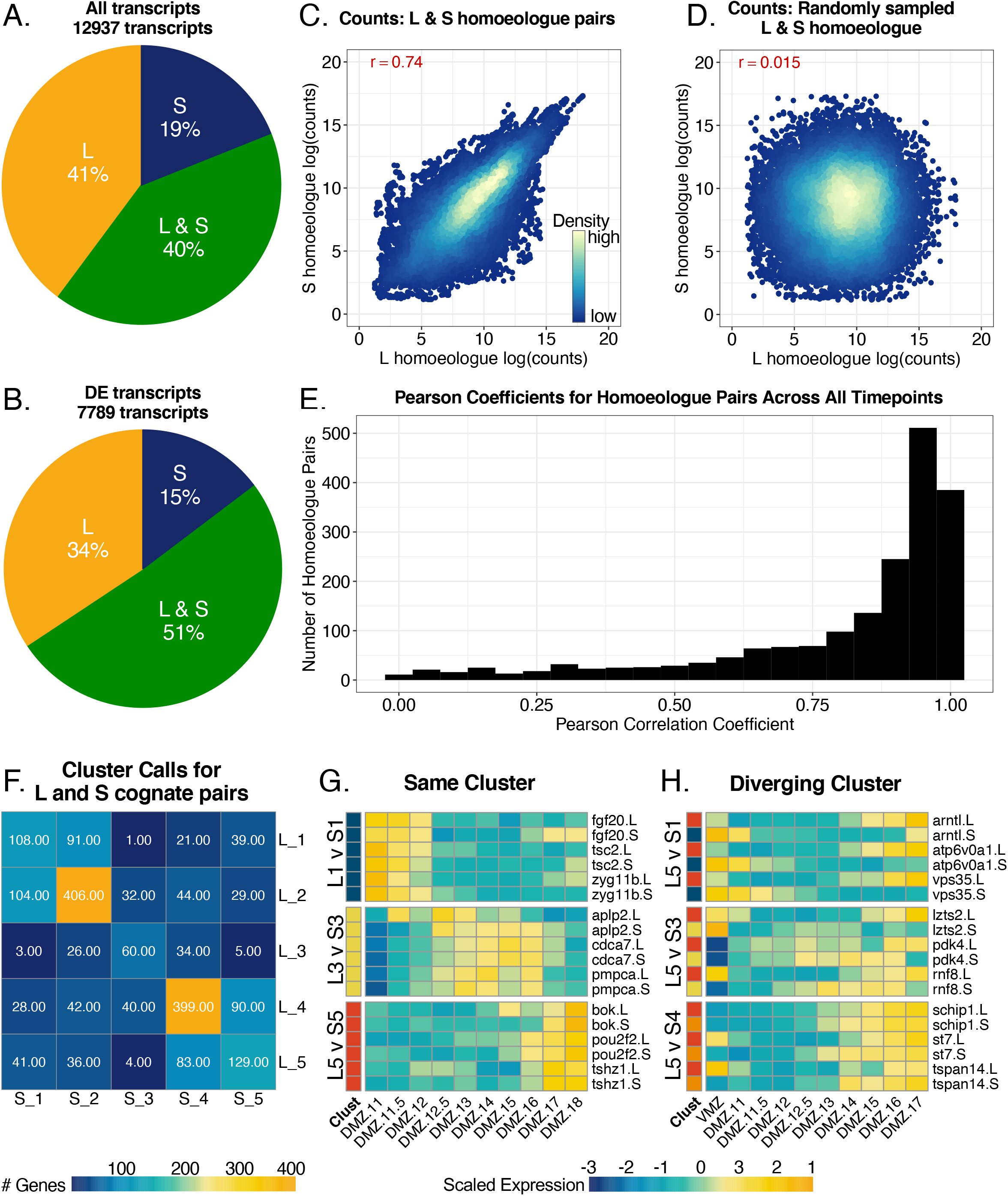
L and S homoeologues are differentially represented and may have differential expression between cognate pairs. A) Pie chart demonstrating the distribution of only L homoeologues, only S homoeologues, and genes with L and S homoeologues across all transcripts in our dataset. B) Distribution of homoeologues in only differentially expressed genes. Genes in the L and S category must have both L and S homoeologues be differentially expressed. C) Correlation plot for log(counts) of L vs. S homoeologues for cognate pairs in all samples. The red number represents the pearson correlation coefficient. D) Correlation of randomly sampled L and S homoeologue log(counts) from all samples. E) Histogram of pearson correlation coefficients calculated for each differentially expressed gene with and L and S homoelogues across DMZ time. F) Heatmap representing the number of genes with L and S homoeologues with each unique cluster call pairing. G) Heatmap showing examples of congruent expression patterns of homoeologues called in the same cluster. H) Heatmap showing examples of expression patterns of homoeologues that were called in different clusters. Color represents expression of gene scaled by row in heatmaps from (G/H).

As a way of visualizing correlation in temporal gene expression patterns between homoeologues, we asked how often the L and S copies of the same gene appeared in the same cluster. The most common observation was that L and S copies were both found in the same cluster, especially the largest clusters, 2 and 4, (Figure 5F), with very similar expression patterns over time. However, cases where the L and S homoeologues were expressed in clusters with opposite temporal dynamics were not uncommon; for example, we found 41 cases in which the L homoeologue was expressed in cluster 5 (early expression) and the S in cluster 1 (late expression). Examples of some of the most tightly correlated and the most strongly anticorrelated homoeologues are shown in Figure 5G,H. This analysis emphasizes that the two *X. laevis* subgenomes are far from identically regulated in DMZ explants. While it has become commonplace in expression analysis to refer to the gene symbol only, our data suggest that tracking the gene expression levels and temporal dynamics of homoeologues is critical to choosing markers with high fidelity to a specific tissue type or temporal pattern.

Finally, the *Xenopus* DMZ has been studied most extensively in the context of collective cell movements termed convergent extension (Huebner and Wallingford, 2018; Wallingford et al., 2002), so we next explored the dynamics of expression for genes implicated in this process. Previous studies of *Drosophila* border cells and migrating cardiac precursors in *Ciona* suggest that transcriptional signatures may be useful for identifying novel players in in the directed migration of cells (Borghese et al., 2006; Christiaen et al., 2008; Morrison et al., 2017; Schwarz et al., 2012; Wang et al., 2006), and our previous exploration of gastrula stage gene expression did reveal novel regulators of morphogenesis (Popov et al., 2017).

Somewhat surprisingly, then, our data from the DMZ revealed no obvious correlation between gene expression patterns and convergent extension cell movements, either temporally or spatially. For example, Planar Cell Polarity (PCP) signaling is the most well-defined regulatory system for vertebrate convergent extension (Butler and Wallingford, 2017; Devenport, 2014), yet examination of a manually curated set of genes revealed no obvious “pan-PCP” signature (Figure 6). In fact, PCP gene expression was not even enriched in early DMZs compared to VMZs. This finding was surprising because DMZs undergo robust convergent extension at these stages while VMZs do not (Keller and Danilchik, 1988).

**Figure 6.**
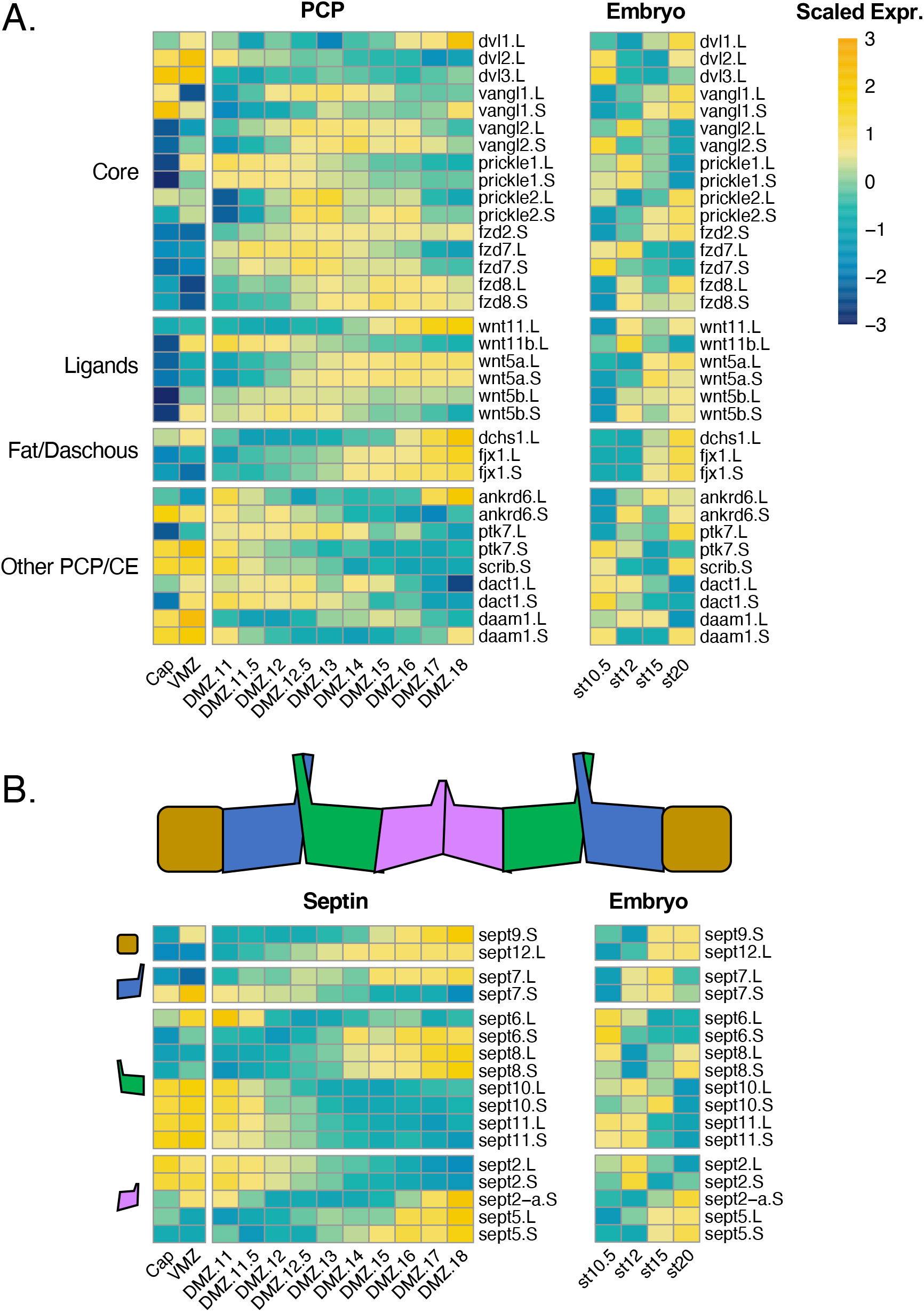
Known PCP pathway genes have little transcriptional concordance. A) Heatmap plotting gene expression of PCP pathway genes in (left) explants and (right) the whole embryo. B) Heatmaps plotting *septin* genes downstream of PCP. Heatmap chunks represent septins representing different protein complex subunits. Color represents expression of gene scaled by row.

Moreover, of the so-called “core PCP” genes (Butler and Wallingford, 2017), only the S form of *prickle1* was robustly differentially expressed in the DMZ at early gastrulation, and all three *dishevelled* genes were actually expressed at higher levels in the VMZ (Figure 6A). Similar variability in expression patterns were observed for both ligands and downstream effectors implicated in convergent extension. Interestingly, however, we observed patterns that suggest a complex relationship between paralogous genes. For example, while Wnt11b was strongly and differentially expressed in the DMZ at gastrula stages, its expression diminished concomitantly with an increase in the expression of the related *wnt11* gene. *Wnt5b/wnt5a* and *fzd7/fzd8* displayed similar reciprocal patterns (Figure 6A).

We next explored the expression of genes encoding proteins that act downstream of PCP signaling in convergent extension. These, too, displayed no perceivable shared signature, though we again observed interesting patterns of expression of functionally related paralogous genes. The septin cytoskeleton provides a good example. Septins have broad roles in actin assembly and cell membrane dynamics and act as obligate hetero-oligomers containing one subunit from each of four different sub-types (Mostowy and Cossart, 2012; Neubauer and Zieger, 2017; Valadares et al., 2017), and we previously showed that Sept2 and Sept7 are required for convergent extension (Kim et al., 2010; Shindo and Wallingford, 2014). Curiously, of the three *septin* 9 sub-group genes, none was highly expressed at early gastrulation, though both *sept9* and *sept12* increased in expression with time (Figure 6B). Of the four genes of the *“septin* 6” sub-family, three were highly expressed at early gastrulation but all decreased over time with the final sub-group member conversely increasing. Among the *septin* 2 sub-family, *sept2* itself was expressed early, but decreased concomitant with an increase in *sept5*.

## Conclusion

Patterns of transcription have long been central to developmental biology, and it was RNA expression cloning that initially discovered the BMP antagonists and Wnt antagonists that give the *Xenopus* DMZ its extraordinary inductive properties (e.g. (Glinka et al., 1998; Sasai et al., 1994; Smith and Harland, 1992)). The dataset reported here provides genome-scale insights into the temporal patterns of gene expression in this tissue when explanted and grown in culture. Our data reveal that these explants develop the full complement of A/P and D/V neuroectodermal patterning, though they do not appear to maintain the D/V pattern over time. The DMZ is equally valuable for its autonomous properties of collective cell motility, and we find that genes controlling cell movement in this tissue are not under the same degree of coordinated transcriptional control as those governing cell fate and signaling. Our data on PCP gene expression are consistent with the high variability of expression seen for other signaling systems throughout *Xenopus* development (Michiue et al., 2017). This result reminds us that mRNA levels are actually very poor predictors of protein levels (Liu et al., 2016; Vogel and Marcotte, 2012), a disconnect that has recently been investigated in *Xenopus* (Peshkin et al., 2015). Thus, while the dynamic proteome of *Xenopus* embryos has been cursorily explored in recent years (Peshkin et al., 2015; Sun et al., 2014; Wühr et al., 2015), our data highlight a critical gap that must be filled by in-depth characterization of the proteome and the protein interactome of *Xenopus* tissues. We also find that overarching trends in the patterns of expression of L and S homoeologues are not adequate to predict which allele should be used when considering molecular markers. Thus, our study provides a useful dataset for continued exploration of an important vertebrate embryonic explant system.

## Acknowledgments

Work in the JBW Lab was supported by the NICHD (R01HD099191) and the NIGMS (R01GM104853). Work in the AEW lab was supported by the NINDS (R01NS099124). TK was supported by the National Research Foundation of Korea (2016R1C1B2009302 and 2018R1A6A1A03025810), the Future-leading Project Research Fund (1.200094.01) of UNIST. TK was also partially supported by the Institute for Basic Science (IBS-R022-D1). KK was supported by the Global Ph.D. Fellowship (GPF) Program through the National Research Foundation of Korea (2017H1A2A1046162).

## Materials and Methods

### Explant preparation, culture and RNA extraction

Fertilized eggs were incubated in 1/3 MMR at 16°C until stage 10.5. 50 DMZ explants were isolated from stage 10.5 embryos using forceps, hair loop, and hair knife, then cultured in 1x Steinberg’s solution. When sibling embryos became stage 11, 11.5, 12, 12.5, 13, 14, 15, 16, 17, and 18, five or six DMZs were collected and dissolved in TRIzol Reagent for each time point. The time interval between the stages was approximately 30 minutes to an hour at room temperature. 20 animal caps and five VMZs were excised from stage 8 and 10.5, respectively. Animal caps and VMZs were dissolved in TRIzol when sibling embryos were at stage 11.

### RNA-Sequencing pre-processing, differential analysis, and clustering analysis

#### Trimming and pseudoalignment

Fastq files were returned from sequencing. Adapter sequences were trimmed from reads and low quality sequences (Phred<33) were removed using Trim Galore!. Kallisto was used for pseudoalignment and counts quantification (Bray 2016). We built a kallisto index from the *X. laevis* transcriptome v9.2. with “kallisto index”. Samples were pseudoaligned and quantified with the “kallisto quant” command. Kallisto quant returned .tsv files for each sample.

#### Counts table

Each sample .tsv file was read into R and compiled to a counts table with the “tximport” package command “tximport” (options: type= “kallisto”, txOut=TRUE) (Soneson 2015). The counts table output was made to be a DESeq object with DESeq2’s command “DESeqDataSetFromTximport” (options: colData= sample metadata, design= ~Replicate +Stage_Tissue), where Stage_Tissue identifies samples according to both the explant type and stage of tissue. The resulting DESeq object was filtered for transcripts with low counts. A log transformed counts table was made for downstream analysis with “rlog” (options: blind=FALSE).

#### Differential expression analysis

##### DESeq2 Likelihood Ratio Test (LRT)

Differential analysis was performed in DESeq2 with the likelihood ratio test across all samples using the command “DESeq” (options: test= “LRT”, full= ~Replicate + Stage_Tissue, reduced= ~Replicate) (Love 2014). Differential analysis results were read with “results” (options: name= “Intercept”, independentFiltering=TRUE, alpha=0.5, pAdjustMethod= “BH”, parallel=TRUE). The p-values were then adjusted by the Benjamini & Hochberg method, and transcripts with an adjusted p-value < 0.001 were kept for clustering. For supplementary figure 1, an additional filtering step was applied to indicate enrichment of DMZ stage 11 over VMZ: logcounts(DMZ)-logcounts(VMZ)>1.

##### DESeq2 Wald Test

For direct pairwise comparison of stage 11 DMZ and VMZ, we used DESeq2’s Wald test with the command “DESeq” (options: test = “Wald”, full= ~Replicate + StageTissue). The differential analysis call was followed by the “lfcShrink” command to shrink dispersion of the logFC calls. The results were filtered to genes with a logFC > 1 and padj < 0.05.

##### STAR/limma

For a secondary analysis type in Supplementary Figure 1 “Limma”, the RNA-Seq samples were aligned with STAR as previously performed in (Session et al., 2016) and enrichment was tested with limma. In limma, the command “eBayes” was used to call differential expression. Results were filtered to genes with a logFC > 1 and padj < 0.05.

#### Sample Clustering and GO analysis

Clustering analysis was performed for only differentially expressed genes (by the LRT method) from the DMZ samples. The log transformed counts for each replicate were averaged, and the resulting matrix of averaged values for each sample were clustered using the package Pheatmap’s command “pheatmap” (options: scale= “row”, kmeans_k = 5, cluster_rows = TRUE, cluster_cols = FALSE). We determined the number of clusters by calculating the sum of squares for k-means clustering of 1-15 clusters, then plotting the number of clusters by the sum of squares variation. Using the heuristic “elbow method” we selected 5 clusters. Gene ontology analysis of clusters was performed by using the list of genes in each cluster as input to gProfileR2 (options: organism = “hsapiens”, sources = “GO:BP”, evcodes = TRUE) (Uku et al 2019). Prior to input, “.L” and “.S” designations were removed from gene names. Returned GO terms were filtered for a term size less than the median term size of all returns to restrict results to the most specific processes.

### Use and manipulation of previously published data

Figure 2 DEG comparison: “Popov” refers to genes recovered from the supplemental table 1 from (Popov et al., 2017). “Ding” refers to genes recovered from supplemental table 3 from (Ding et al., 2017)

Figure 3/4/6 Whole Embryo gene expression: The “Embryo” heatmaps refer to genes expression measured in (Session et al., 2016). The updated counts table (aligned with genome 9.2) was retrieved from Xenbase. The counts table was read into DESeq for normalization with “DESeqDataSetFromMatrix”. The counts were normalized with “rlog” (options: blind=FALSE).

### Data deposition

Raw RNA-Seq data will be deposited on GEO upon acceptance of the paper.

---

***Supplementary Figure 1.** Variation in differential expression analysis pipelines can lead to low recall in differentially expressed genes*. A) Venn diagram showing the overlap in the number of gene called enriched in DMZ stage 11 samples over VMZ samples. Box below venn diagram indicates main differences in differential expression analysis for each approach. B) Heatmap showing expression of 100 union genes between analysis workflows. Color represents expression of gene scaled by row.

***Supplemental Table 1.*** This table contains the DESeq2, rlog transformed counts for all transcripts sequenced in our dataset. We include columns indicating if the genes were differentially expressed by the likelihood ratio test (“DE”), the adjusted p-value for the DE designation (“padj”), and the cluster each DE gene was called in (“Cluster”).

